# A Theta Band Network Involving Prefrontal Cortex Unique to Human Episodic Memory

**DOI:** 10.1101/140251

**Authors:** James J Young, Peter H Rudebeck, Lara V Marcuse, Madeline C Fields, Ji Yeoun Yoo, Fedor Panov, Saadi Ghatan, Arash Fazl, Sarah Mandelbaum, Mark G Baxter

## Abstract

Episodic memory, everyday memory for events, is fundamental to our lives. We tested patients undergoing intracranial electroencephalography (intracranial EEG) monitoring for the localization of medically-refractory epilepsy on a well-characterized paradigm that requires episodic memory. Here we report that an anatomically diffuse network characterized by theta-band (4-7 Hz) coherence is activated at the time of a choice that requires episodic memory. This distinct network pattern of oscillatory activity is absent in situations that do not require episodic memory. The episodic network we identified spans prefrontal and temporal lobes and coherence between these areas was greatest during memory encoding. Our data provide novel empirical evidence for a set of brain areas that supports episodic memory in humans.

**Significance Statement:** Synchronous oscillations of electrical fields are thought to support brain processes such as language, movement, memory and learning. However, little is known about which types of synchronous oscillations are needed for which process. We performed invasive recordings of electrical activity in patients undergoing surgical management for epilepsy while they perform different types of memory tasks. We found synchronous oscillations in the theta band (4-7 Hz) correlate with performance in an episodic memory. This pattern of oscillations activates a distinct network of brain regions from a task that does not require episodic memory. This finding advances our understanding of the brain systems that support each process.

## Introduction

Memory for every day events – known as episodic memory – is integral to our daily lives and is critically affected in dementia and other neurological disorders. Although much is known about the parts of the brain involved in forming, storing, and retrieving episodic memories, much less is known about the patterns of neural activity that support this process in humans. One hypothesis derived from studies in animal models is that memories are formed and retrieved through dynamic interaction between distributed brain areas. Specifically, synchronous oscillations in electrical activity are thought to support mnemonic function by binding information from spatially-distinct parts of the brain into a functional network [1–4]. However, empirical support for this hypothesis in humans is scarce [5], particularly in the setting of invasive recordings which offer superior spatial resolution to scalp EEG and magnetoencephalography. While some studies have examined synchronous oscillations in the setting of episodic memory [6,7] with invasive recordings, none have compared episodic and non-episodic memory to identify the types of oscillatory synchrony critical for each task. Accordingly, determining whether synchronous oscillations support human episodic memory would provide vital insight into this fundamental aspect of human cognition.

Here we sought direct empirical support for this hypothesis by testing 7 patients undergoing intracranial EEG on episodic and non-episodic memory tasks. Recordings of neural activity were made while patients were tested on an object-in-place scene memory task implemented on an iPad, allowing straightforward administration in the hospital setting. In the object-in-place scene memory task, multiple pairs of fractal visual targets are depicted on a colored background, representing a complex episode which includes target, location, contextual cues and time. Rapid learning of object-in-place scene discrimination problems assess episodic memory [8–10] and requires interaction between frontal and temporal cortex [11–13]. By contrast, replacing the scene component of the discriminations with a white background eliminates the contextual cues surrounding the targets. This results in memory being supported by non-episodic memory systems and removes the requirement for interaction of frontal and temporal cortex for normal performance [11,12]. The episodic memory task allows the subject to associate the target with nearby contextual information, but this information cannot be utilized in the non-episodic memory task. By testing patients with intracranial electrodes while performing episodic and non-episodic tasks, we demonstrate an anatomically diffuse theta band coherence network that appears to support episodic but not non-episodic memory in humans. Further, we provide novel empirical support in humans for the hypothesis that synchronous oscillations coordinate activity between brain regions and form them into a functional network.

## Activity During Performance of Episodic and Non-episodic Memory Tasks

We recorded activity from 854 locations in the 7 subjects during performance of episodic and non-episodic memory tasks (Figure 1, Supplementary Table 1). Of these, 259 showed interictal discharges and 61 were located in the seizure onset zone (as determined by a board-certified epilpetologist), and were excluded from further analysis. An additional 13 electrodes in the left hemisphere and 15 in the right hemisphere were excluded because they were located in white matter. Of the remaining 506 locations, 74 were in the left hemisphere and 432 in the right hemisphere (Supplementary Figure 1). The raw spectral power for each electrode was calculated as well as the pairwise coherence between each electrode and every other electrode for the delta [1,4) Hz, theta [4,8) Hz, alpha [8,15), and beta [15,30) bands. Spectral power and coherence were compared between episodic and non-episodic memory tasks to identify specific oscillatory signatures of episodic memory formation and retrieval.

**Figure 1:**
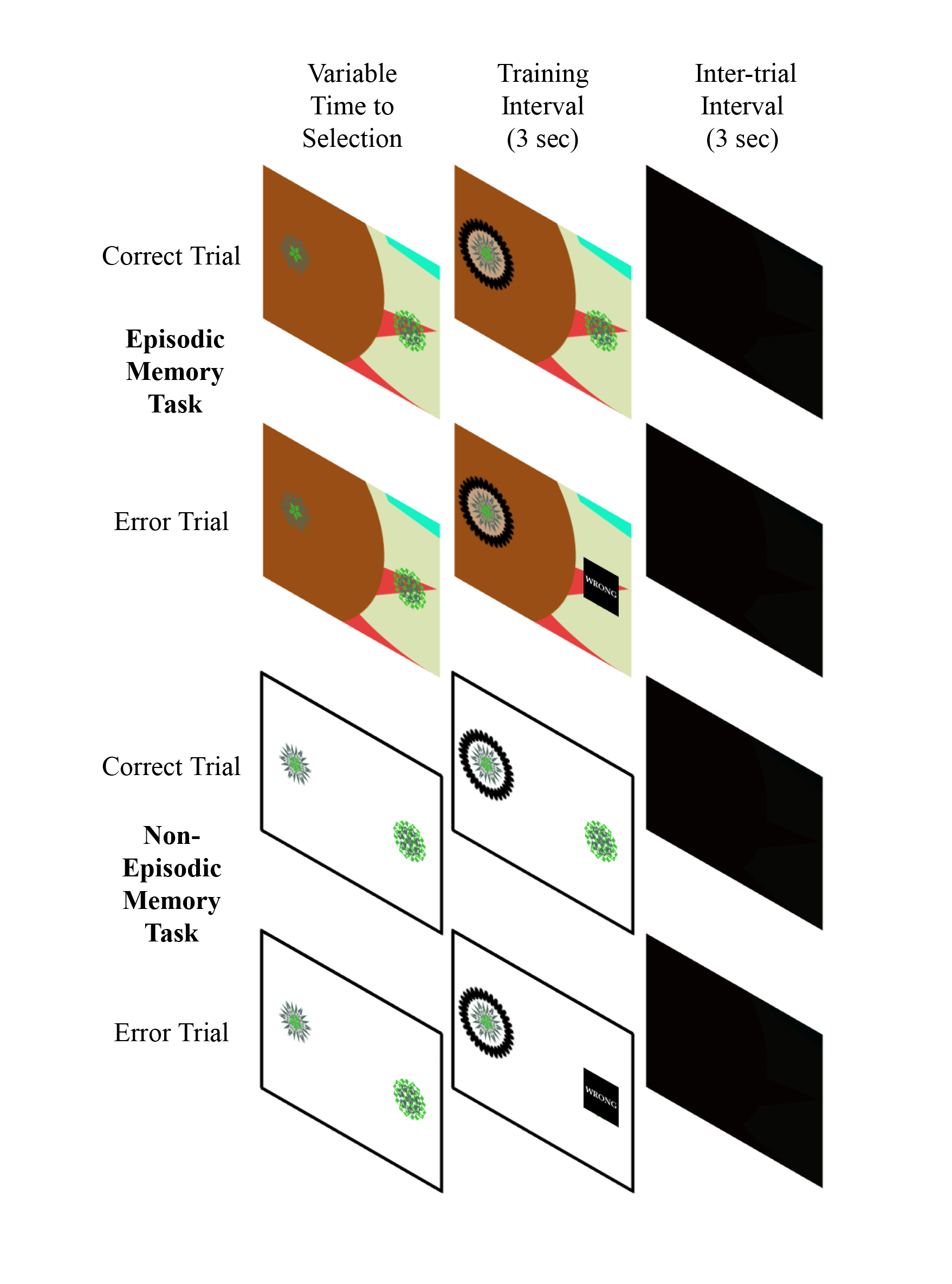
In the episodic memory task, 2 fractal target images are presented on a colored background filled with randomly populated geometric figures. If the subject selects the correct target, the target flashes for 3 seconds, followed by a 3 second Inter-trial Interval (ITI) before the next trial. If the subject selects the wrong target, the wrong target is replaced with a black square that says “WRONG” and the correct target flashes for 3 seconds, followed by a 3 second ITI. There is a 16 trial block where the subject has never seen the exemplars before. This 16 trial block is then repeated 4 times for a total of 80 trials. The non-episodic memory task differs from the episodic memory task only in that the background is white rather than colored.

To reduce the chance of false positives, we defined the threshold for statistical significance in four ways. First, for an effect to be marked as significant, there had to be a statistically significant difference in activity (by Bonferroni-corrected Student’s t-test) between the episodic and non-episodic tasks for at least 100 ms contiguously. Second, each effect had to be present and statistically significant in at least two of the seven subjects individually. Third, any effects had to correlate with performance (by Bonferroni-corrected Pearson’s R) during the time of significant difference in activity between the tasks. Fourth, any effects had to occur at a point where the activity had a significant imaginary component of complex coherence – a part of the wavelet used to extract the oscillations that ignores activity occurring at a zero time lag and thereby excluding effects of volume conduction [14].

## Theta Band Coherence Increases Around the Choice Point in the Episodic Memory Task

Applying these criteria, there was a significant and broadly anatomically distributed network of pairwise theta-band coherence that occurred exactly or very closely after the choice point and was closely related to behavioral performance in the episodic memory task (Figure 2A, 2C). This was specific to the episodic memory task, and not observed in other frequency bands. Further, the increase in theta band pairwise coherence is not observed in the non-episodic memory task. Note that the effect is most distinct in the interaction between the frontal and temporal (medial and lateral) electrodes, but also includes interactions with the parietal and occipital cortices. Although the alpha and beta frequency bands showed similar patterns, none were modulated to the same degree or showed such a tight relation to behavior (Supplementary Figure 1). There were no significant effects observed in the delta band. There is an increase in overall theta band power, particularly in the mesial temporal region, but there is no difference in theta band power between the tasks (Supplementary Figure 2). Further, this effect occurs when the imaginary part of coherence – coherence modified to ignore activity occurring at a zero time lag – also shows a statistically significant increase (Supplementary Figure 3) indicating that the effect is not from volume conduction.

**Figure 2:**
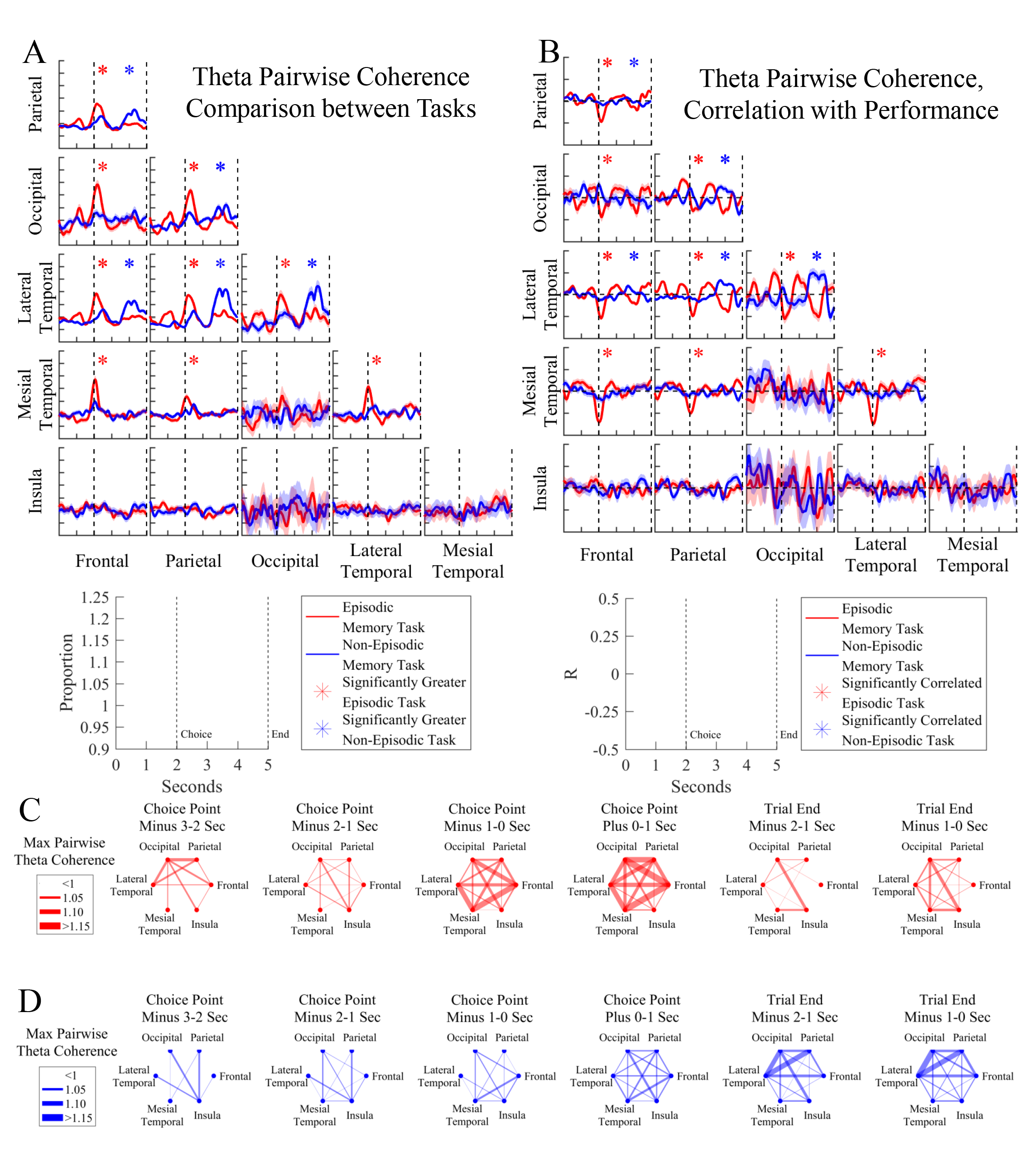
(A) The proportion change in pairwise theta coherence between the two indicated brain regions (row and column labels) during the performance of the episodic memory (red) and non-episodic memory (blue) tasks. The first vertical dotted line indicates the choice point. The second vertical dotted line indicates the end of the trial. A red or blue asterisk indicates a significant difference in theta coherence between the tasks (Bonferroni corrected p < 0.05 by Student’s t-test). (B) The correlation between pairwise theta coherence between the two indicated brain regions (row and column labels) and percent correct during that trial block for the episodic memory (red) and non-episodic memory (blue) tasks. An red or blue asterisk indicates a significant correlation between theta coherence and performance (Bonferroni corrected p < 0.05 by Pearson’s R). (C) Maximum pairwise theta-coherence between the connected brain regions during the listed interval for the episodic memory task. (D) Maximum pairwise theta-coherence between the connected brain regions during the listed interval for the non-episodic memory task.

This anatomically broadly distributed network shows a negative correlation with trial block performance at the time of the choice (Figure 2B). This indicates that early trial blocks where the performance was lower show a larger increase in theta pairwise coherence than later blocks where performance was higher, i.e. early in blocks where the subject must encode new exemplars, there is the greatest increase in pairwise theta coherence in this network. The largest and sharpest negative correlations are observed between the frontal lobe and the temporal cortices, but the negative correlation with activity recruits the parietal and occipital lobes as well.

In an effort to more accurately anatomically localize this theta band coherence network, analogous comparisons were constructed for pairwise coherence between all the regions in the DKT40 Atlas [15] as well as the hippocampus and amygdala. The peak height of the increase in theta band coherence during the episodic task was calculated for the interval between 1 second before and 1 second after the choice. This is indicated in Figure 3. The maximum theta coherence increase between each brain region is shown. The areas with maximal increase involved in this network include medial orbitofrontal, pars opercularis, superior frontal, rostral anterior cingulate, precuneus, inferior temporal, and parahippocampal gyri along with the hippocampus and the amygdala. Similarly, the network in which the increase in pairwise theta coherence was the greatest also corresponds to the greatest negative correlations with performance (Figure 4).

**Figure 3:**
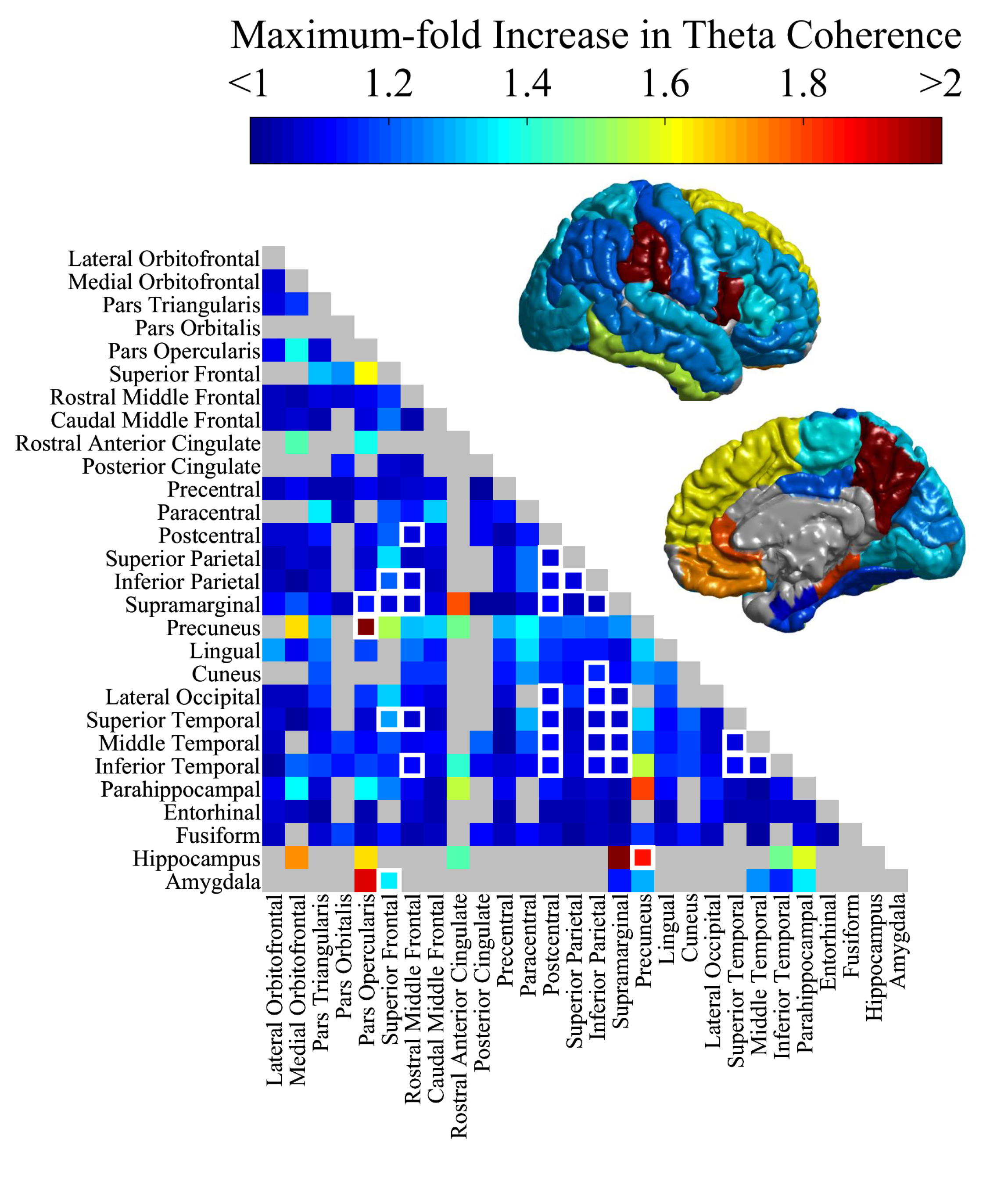
The lower triangle indicates the peak height of the increase in theta band pairwise coherence between the brain regions listed (row and column) in the episodic memory task during the period between one second before and one second after the choice. The color bar above the figure indicates the proportion change. White boxes surround connections that show a statistically significant difference between the episodic and non-episodic memory tasks. Gray boxes indicate connections for which there was no data. The maximal increase in pairwise theta coherence between the depicted region and other parts of the network around the choice point during performance of the episodic memory task is depicted on the brains.

**Figure 4:**
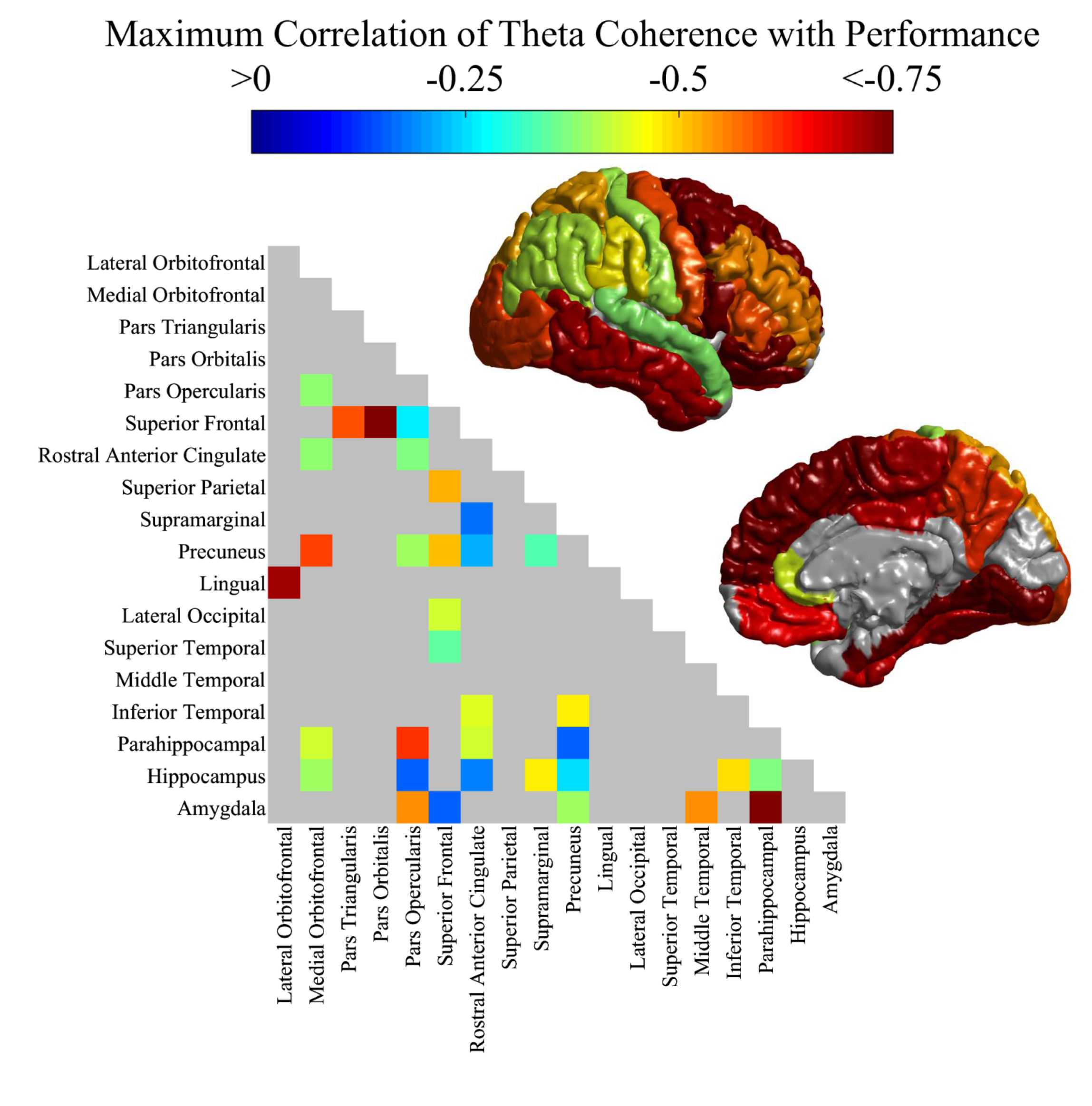
The lower triangle indicates the maximum depth of the negative correlation with performance and pairwise coherence between the brain regions listed (row and column) in the episodic memory task during the period between one second before and one second after the choice. For clarity only connections with an increase of 1.25-fold or greater are depicted. The color bar above the figure indicates the correlation value. Warm colors indicate a stronger negative correlation. The maximal negative correlation coefficient between the depicted region and other parts of the network around the choice point during performance of the episodic memory task is depicted on the brains.

## Theta Band Coherence Increases Around the Trial End in the Non-Episodic Memory Task

Comparison of the episodic and non-episodic memory tasks showed that a theta band network is activated around the choice point in the episodic but not the non-episodic memory task. However, theta band activation does occur at a different point in the trial during the non-episodic memory task. Figure 2A and 2D also show significant increases in pairwise theta band coherence between the frontal, parietal, occipital and lateral temporal lobes. This effect occurs during the last two seconds of the trial – following the choice and immediately prior to the intertrial interval. Further, the size of the theta band coherence increase in the non-episodic memory task is positively correlated with performance (Figure 2B). There is a statistically significant increase in theta band power in the frontal lobe during this period, and a trend toward significance in the other included regions (Supplementary Figure 2). However, the imaginary part of coherence is also greater than baseline during this period (Supplementary Figure 3), suggesting that the effect is not related to volume conduction.

To further isolate anatomical specificity of this effect, the peak height of the increase in theta band coherence during the non-episodic memory task was calculated for the interval of the 2 seconds at the end of the trial. The maximum theta coherence increase between each brain region is shown. Figure 5 shows an overlapping but different network is activated during the control task including, medial orbitofrontal, pars orbitalis, superior frontal, postcentral, inferior parietal, supramarginal, lateral occipital, entorhinal and fusiform cortex. Hippocampus and amygdala are much less involved than the version of the task requiring episodic memory. These brain regions correspond to the areas of maximal positive correlation with performance (Figure 6). Thus, analysis of the pairwise theta coherence during the version of the task that does not require episodic memory shows an independent network, activated late in the trial that positively correlates with performance.

**Figure 5:**
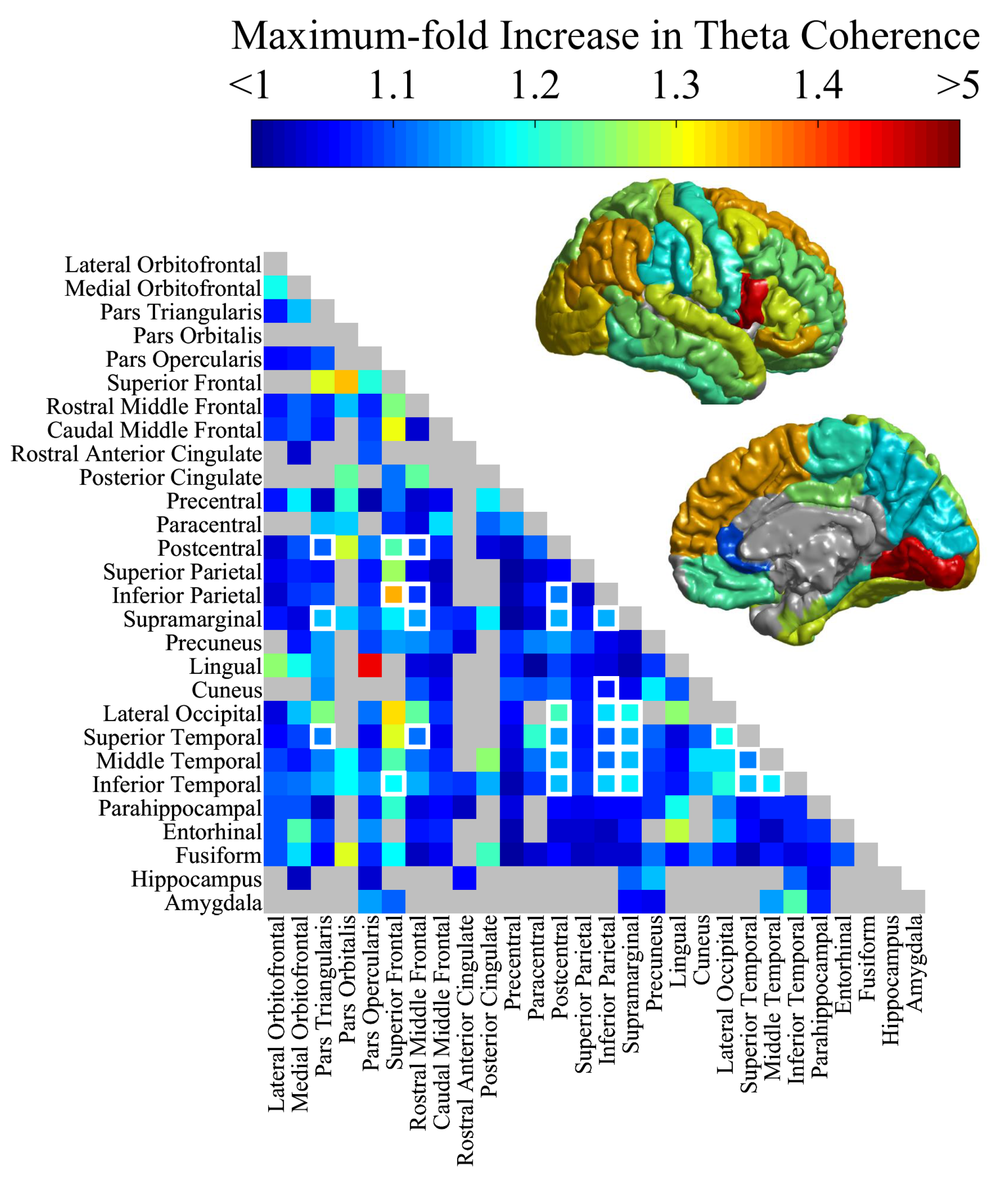
The lower triangle indicates the peak height of the increase in theta band pairwise coherence between the brain regions listed (row and column) in the non-episodic memory task during the last two seconds of the trial. The color bar above the figure indicates the proportion change. White boxes surround connections that show a statistically significant difference between the episodic and non-episodic memory tasks. Gray boxes indicate connections for which there was no data. The maximal increase in pairwise theta coherence between the depicted region and other parts of the network around the trial end during performance of the non-episodic memory task is depicted on the brains.

**Figure 6:**
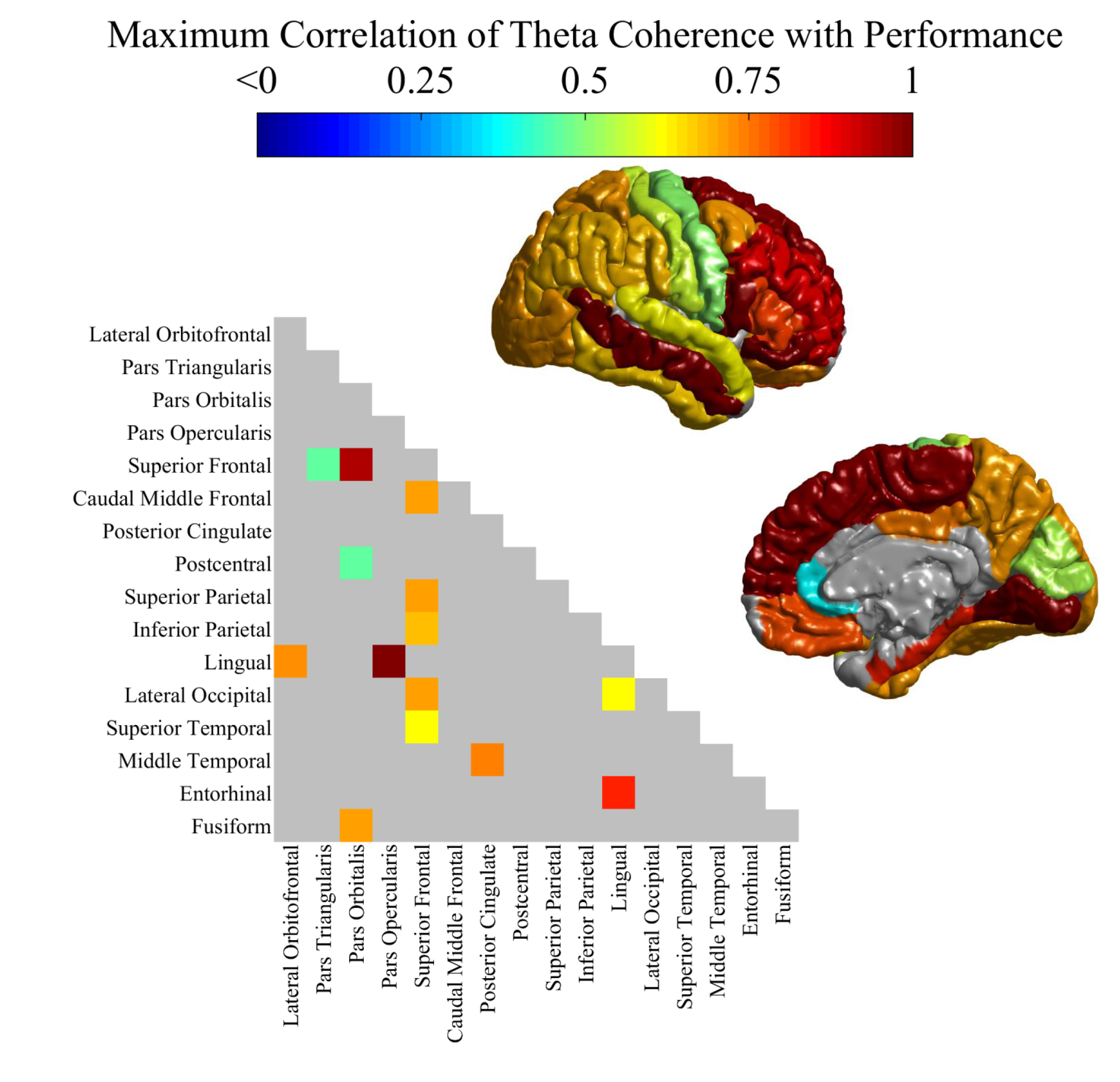
The lower triangle indicates the maximum positive correlation with performance and pairwise coherence between the brain regions listed (row and column) in the non-episodic memory task during the last two seconds of the trial. For clarity only connections with an increase of 1.25-fold or greater are depicted. The color bar above the figure indicates the correlation value. Warm colors indicate a stronger positive correlation. The maximal positive correlation coefficient between the depicted region and other parts of the network around the trial end during performance of the non-episodic memory task is depicted on the brains.

## Discussion

We examined the oscillatory activity and synchrony during the performance of an episodic memory task versus a non-episodic memory task in 7 patients undergoing intracranial EEG for surgical management of epilepsy. Based on synchrony in the theta band, we identify an anatomically broad network including the frontal, temporal and parietal lobes (Figure 2). This network shows activation in theta coherence at the choice point during the episodic memory task, and the degree of activation negatively correlates with task performance. This suggests increased theta coherence may be related to a learning signal to help consolidate the episode for later retrieval. The anatomic network identified here conforms to lesion data in non-human primates and humans [10–12,16,17] and with the anatomical connectivity between the prefrontal cortex and the hippocampus and amygdala [18]. It is also consistent with the results of fMRI experiments [19] and scalp recordings [20–22] during episodic memory tasks which show a broad network including elements of the mesial temporal, prefrontal and parietal cortices. The inclusion of parts of the parietal lobe, particularly the precuneus, which has produced conflicting results in imaging studies during episodic memory [19,23] but has been previously shown to participate in a synchronous network during episodic memory [6], is consistent with a role for the parietal lobe in episodic memory retrieval.

An overlapping but different network including parts of the temporal and parietal lobes was activated during the performance of the non-episodic memory task (Figures 2 and 3). This network positively correlated with performance. We note that few of these brain regions outside the temporal lobe have been implicated by lesion studies in the performance of the non-episodic memory task. We hypothesize that this effect may be related to activation of an attentional network that supports but is not required for performance of non-episodic tasks. It could also reflect greater engagement of episodic memory systems once object-reward associations have been acquired by subcortical, “habit” memory systems [24]. Whatever the functional significance of this network, the differentiation in anatomical components, timing relative to stimulus onset, and association with task performance indicates its distinctiveness from the network that is triggered by episodic memory.

Our results build on prior human intracranial recordings that have highlighted a role for synchronous oscillations in episodic memory [6,7]. For instance, Watrous et al.[6] show successful memory retrieval – using either spatial or temporal cues – increases 1-10 Hz phase synchrony diffusely throughout the brain with a hub in the parahippocampal gyrus. For the retrieval of place information, phase synchrony was highest at 1-4 Hz, where as it was highest in the 7-10 Hz range for information about time of retrieval, and this synchronization differentiated successful and unsuccessful retrieval. Here, by including hippocampal electrodes and a non-episodic memory task, we show that very different networks are recruited during episodic and non-episodic processes. Notably, we found that during episodic memory, activity is divergent between the entorhinal cortex and the parahippocampal gyrus and between the hippocampus and entorhinal cortex. Similarly, Burke et al.[7] show in a word encoding and retrieval task that theta phase synchrony increases between the (predominantly left) temporal and the prefrontal lobes, followed by a prolonged and large decrease. These synchrony changes are much more prominent in successful than unsuccessful encoding. However, they also did not include a non-episodic comparison task. By doing so in this study, we were able to compare the networks involved in the two tasks and relate our findings to lesion data about the brain regions required for task performance. Finally, these results extend previous reports that have examined phase amplitude coupling and phase coding in the setting of episodic memory [25], perceptual categorization [26] and contextual responding [27], all of which show the importance of these nested oscillations in episodic memory encoding, but did not analyze network participation at distant sites.

Our findings also provide unique insight into the how different synchronous networks are recruited in two neuropsychologically well-characterized behavioral tasks that require different forms of memory. The finding that the theta band coherence network includes the sites that have been demonstrated in lesion studies to be required for task performance does not prove that theta coherence is required for episodic memory, but it does add additional weight to this hypothesis. Further it provides a clear time course for the requirement for synchronous oscillation and may represent the neurophysiologic basis through which diverse brain regions are recruited into a functional network.

It is important to address possible confounds to these results including the effect of volume conduction and activity caused by motion of the arm. It is unlikely that these results are secondary to volume conduction for two reasons. First, intervening brain regions between the frontal and temporal lobes such as the insular cortex do not show theta band coherence, contrary to what would be expected if the effect was due to volume conduction. Second, analysis of the imaginary part of coherence removes the presence of synchronous activity at a zero time lag, and the activity remains different from baseline in this analysis. Likewise, it is unlikely that the observable effect is because of motion of the arm towards the target for several reasons. First, the motion is essentially identical in the episodic and non-episodic memory task, yet the two tasks activate very different networks. Second, the effect in the non-episodic memory task appears at the trial end, after motion has ceased. Third, the effect appears in both hemispheres, regardless of the arm that was used to make the choice.

In addition, several aspects of these experiments suggest caution in over-generalizing these results. First, because of limitations in the sampling rate of the behavioral output (behavioral flags were determined manually on the clinical video -- 30 fps), we did not attempt to evaluate effects that occurred at frequencies greater than 30 Hz, which are likely functionally significant [7,27–29]. Second, brain coverage with electrodes was dictated by clinical necessity. There was relatively less coverage of the left hemisphere as compared to the right because non dominant lesions are more eligible for resective surgery. As a consequence, we could not determine whether observed effects were more prominent in the left or the right hemisphere.

An influential theory posits that that synchronous oscillations support cognitive functions by binding information from distant parts of the brain into a functional network [1–4]. Here we show that episodic memory is associated with synchronous activity in a network comprising distant brain regions in frontal, temporal, and parietal lobes. The degree of activity in this network correlates with the accuracy of choices dependent on episodic memory, and choices based on non-episodic memory processes do not require the same network. The size of the network is much larger than what would be suggested by lesion studies, and more closely matches the results of fMRI [19]. Thus, our data provide empirical support for the hypothesis that synchronous activity binds together diverse brain areas into distinct functional networks to support cognition in humans.

## Methods

### Subjects

7 patients undergoing surgical management for treatment of medically refractory epilepsy were consented for participation while undergoing intracranial EEG. The study was conducted according to the principles of the Declaration of Helsinki, and the consent documentation and procedure were approved by the Mount Sinai Hospital Institutional Review Board (IRB). Descriptive information about the subjects is in Supplementary Table 1.

### Task

The subjects were tested using an implementation of the episodic and non-episodic memory tasks on a iPad 2 (Apple, Inc., Cupertino, CA). The iPad was placed before the subject on each testing day at eye level at approximately 20-30 cm away. Environmental distractors were minimized during testing by closing the patient’s room door and preventing them from talking. The task involves the presentation of a two colored fractal target images [30] presented on colored background with randomly generated geometric figures (episodic memory task) or a white background (non-episodic memory task) (Figure 1). Evidence from lesion studies in humans and non-human primates suggest that an intact prefrontal cortex, medial temporal lobe, and interaction between the two, are required for performance of the episodic memory but not the non-episodic memory task [11]. The same two colors were used to generate the two target fractals, and these colors were balanced against to the background to maximize discriminability for the Scenes task. The size of the targets was 100 x 100 pixels on a screen size of 1024 x 768 pixels. Targets were randomly distributed on the screen such that at least 700 pixels separated their centers, and the location of both targets was maintained throughout testing for each target.

During each trial, 2 fractal target images are presented on the background from the beginning of the trial. The subject then selects one of the two targets. If the subject selects the correct target, the target flashes for 3 seconds, followed by a 3 second Inter-trial Interval (ITI) before the next trial. If the subject selects the wrong target, the wrong target is replaced with a black square that says “WRONG” and the correct target flashes for 3 seconds, followed by a 3 second ITI. There is an initial 16 trial block where the subject has never seen the exemplars before. Performance during the initial presentation block was balanced such that the subject would necessarily get 50% of the trials correct, and their choices were propagated throughout the rest of testing. The 16 trial block was then repeated 4 times for a total of 80 trials.

### Data Collection and Pre-Analysis

Electrophysiological data was collected for all subjects using the clinical equipment, a XLTEK 128 EMU headbox connected to the Natus XLTEK EMU40 amplifier (Natus Medical Incorporated, Pleasanton, CA). Sampling rates were between 500 and 1028 Hz. The behavioral flags were correlated with the recording data using the Natus Database software. The three behavioral flags were the time of trial start, the choice point, and the end of the trial. These were manually placed in the recording files by reviewing the associated video recording (sampled at 30 fps). The resultant files were pruned and parsed into trials based on the behavioral flags. The video of testing was reviewed, and trials were the subject was distracted, talked during the trial, or waited greater than 10 seconds before making a selection were excluded from further analyses. All pre-analysis and analysis was performed on MATLAB (Mathworks, Natick, MA) using the FieldTrip software library [31]. Recordings for each trial were referenced to the moving average of all electrodes. Each trace was locally detrended. 60 Hz noise was removed using a notch filter.

### Electrode Localization

Localization of electrodes was performed using pre-operative MRIs and post-operative CTs. Coregistration of MRI and CT was performed using FreeSurfer (http://surfer.nmr.mgh.harvard.edu/), and the location of each electrode was selected on the postoperative CT. A parcellated image of the patient’s cortical surface was generated using FreeSurfer from the T1 series of the pre-operative MRI. Cortical parcellation is by the DKT40 Atlas ^27^. The position of the electrodes was then projected onto the cortical surface, and the brain region identified was used as each electrodes location. Confirmation of correct location was performed using images taken during the surgery. The report from the recording was reviewed, and those electrodes that showed interictal discharges or those within the seizure onset zone were excluded from all subsequent analyses.

### Analysis

Because the behavioral flags were established using the video recording of the subject performing the task, analysis focused on frequency bands lower than 30 Hz, because of concern that averaging due to inexact behavioral flag placement would obscure relevant effects. Consequently all trial traces were downsampled to 100 Hz. Spectral powers were calculated using the FieldTrip using a complex Morlet wavelet transformation (wavelet width = 7). Spectral power was calculated for all electrodes from between 1 to 30 Hz, calculated at 1 Hz increments. Pairwise coherence between each electrode and every other electrode was calculated using a complex Morlet wavelet for delta [1,4) Hz, theta [4,8) Hz, alpha [8,15), and beta [15,30) bands. Subsequently, coherence refers to the modulus of complex coherence from this analysis. Where specified, the term imaginary part of coherence refers to the imaginary part of complex coherence from the Morlet wavelet transformation, an analysis which isolates those parts of coherence that occur at a non-zero phase lag, excluding the possibility of contamination with volume conduction [14]. Average spectral power and coherence was calculated for each subject for each electrode (or pair of electrodes) for each 16 trial block.

### Statistical Analysis

The results of calculations for spectral power – grouped by frequency band -- and coherence between each electrode at each frequency band were grouped according to the anatomical location of the electrodes. Large grouping including the lobe in which each electrode was located and smaller grouping – divided by the cortical parcellation and subcortical segmentation of the subject’s brain imaging. Each measure was subjected to three statistical tests. First, the coherence or spectral power at a particular band (divided by the mean at that band for that trial block) was calculated for each block and anatomic location/pairing. At each time point, whether the activity was different between the episodic and non-episodic memory tasks for each time point (a 1/100 second bin) was evaluated using a Student’s t-test. This was calculation was synced to two time points (trial start and choice point) for all the subjects combined and for each subject individually. Second, the coherence and spectral power at a particular band (divided by the mean at that band for that trial block) was correlated with performance during that block using Pearson’s R. (The first block was excluded because the subjects had never seen the exemplars during this block.) This was calculated synced each time point (trial start and choice point) in all time bins (a 1/100 second bin) for all the subjects combined and for each subject individually. Finally, the imaginary part of coherence, which isolates those parts of coherence that occur at a non-zero phase lag [14], was calculated synced each time point (trial start and choice point) in all time bins (a 1/100 second bin) for all the subjects combined and for each subject individually. This measure applies only to the coherence calculations, not to the spectral power. The results of this calculation were tested to determine whether at each time point, the activity differed from 1 – indicating the baseline imaginary coherence between regions at that band – to determine whether changes in activity occur at a non-zero time lag. The results of all statistical tests were corrected for multiple comparison’s using Bonferroni correction. The total number of statistical tests was 4,301,328, so the p-value criterion was < 1.16E-8. The total number of statistical tests of each type are listed in Supplementary Table 2.

## Acknowledgements

The authors would like to acknowledge Dr. Brad Voytek, PhD for comments on an earlier draft. We would like to acknowledge Dr. Matthew Shapiro for the use of computational resources. Finally, we would like to thank the Leon Levy Foundation and the NINDS R25 (NS8440304) for continuing financial support.

